# MOTLAB: A Weighted Multi-Omics Transfer Learning Framework for Reducing Racial Disparities in Breast Cancer

**DOI:** 10.64898/2025.12.22.696111

**Authors:** Minjeong Baek, Lusheng Li, Vimla Band, Jieqiong Wang, Shibiao Wan

## Abstract

Breast cancer mortality remains markedly higher among Black women in the United States, while the limited representation of Black women in genomic datasets constrains the development of reliable clinical outcome prediction models for this racial group. Conventional transfer learning improves prediction in data-minority groups, but it does not simultaneously optimize omics-specific contributions in multi-omics integration and enhance learning from limited target-domain data. We developed MOTLAB to address both challenges through optimized multi-omics weighting, and data augmentation. In 1,085 female TCGA breast cancer cases, MOTLAB yielded the highest AUROC with three-omics integration across all progression-free interval time settings, reduced prediction error, and lowered calibration error in most settings compared with transfer learning alone. MOTLAB performance remained robust across molecular and clinical subgroups, while MOTLAB-derived risk scores captured breast cancer-relevant pathway, miRNA, and methylation signals. MOTLAB provides an integrated strategy that combines omics-specific weighting, transfer learning, and data augmentation to improve breast cancer clinical outcome prediction in data-minority groups.

## Introduction

Breast cancer (BC) is the most commonly diagnosed cancer among women in the United States, and the second leading cause of cancer death after lung cancer^1^. Notably, BC mortality rate among Black women is 38% higher than that of White women^2^. This difference reflects a persistent racial disparity in BC mortality^3,4,5^. Moreover, social determinants of health (SDOH), including socioeconomic disadvantage and geographic barriers to care, can contribute to disparities in BC treatment and outcomes^6,7,8^. To reduce cancer health disparities, patient navigation, culturally tailored interventions, and community-based programs have been implemented. Patient navigation helps overcome logistical, linguistic, and financial barriers and increase BC screening in groups affected by health disparities^9,10^. Culturally tailored navigation and community-based interventions also increase BC screening in underserved groups^11,12,13^.

Although these approaches are important for reducing BC health disparities, racial minority groups are underrepresented in molecular datasets used for improving precision oncology and predictive modeling.

A central challenge for developing ML approaches is the substantial imbalance in molecular data availability across ancestry groups^14,15,16^. In TCGA, for example, individuals of European ancestry comprise approximately 80% of the cohort, whereas individuals of African ancestry comprise less than 10%^14^. Difference in sample size and data distributions across ancestry groups directly influence the amount of information available for model development and conventional learning strategies yield low predictive performance in data-minority groups^14^. This limitation motivates transfer learning (TL) approaches that leverage knowledge from data-majority groups to improve prediction in data-minority groups^14,17,18^. Moreover, target-domain data augmentation (DA) has been applied to improve predictive performance in other settings^19,20^, while multi-omics data integration capture interdependent information across multi-omics data that is not available from single omics data only and has been reported to improve cancer outcome prediction^21,22,23^. We hypothesized that individual omics modalities differently contribute to prediction and therefore optimized modality-specific weights to consider different contributions of multi-omics data and improve multi-omics integration. Overall, these considerations motivate an integrated strategy that combines TL, DA, and optimized multi-omics integration to improve BC outcome prediction in data-minority group in this study.

To address these limitations, we developed MOTLAB, a weighted multi-omics TL framework designed to improve BC outcome prediction in data-minority groups. MOTLAB integrates mRNA, miRNA, and DNA methylation data by optimizing modality-specific weights of patient–patient similarity representations and incorporates synthetic DA to mitigate data imbalance between data-majority and data-minority groups. The framework transfers knowledge from the data-majority European American (EA) group to the data-minority African American (AA) group using Classification and Contrastive Semantic Alignment (CCSA), a supervised domain adaptation method^24^. In this study, MOTLAB demonstrated the impact of DA and optimized multi-omics integration beyond conventional TL, while maintaining predictive robustness across clinical and molecular subgroups and yielding biologically relevant risk estimates. Thus, these findings support MOTLAB as a strategy for leveraging molecular information from data-majority groups to improve BC outcome prediction in data-minority groups.

## Results

### 2.1. MOTLAB integrated multi-omics, transfer learning, and data augmentation for clinical prediction to mitigate racial disparities

To address racial disparities in BC outcome prediction associated with data imbalance, we developed MOTLAB, a weighted multi-omics TL framework composed of four major steps (**Fig. 1**). We derived the study cohort from TCGA-BRCA by restricting the analysis to female patients and retaining EA serving as the data-majority group and AA as the data-minority group while defining binary progression-free interval (PFI) prediction tasks at 2, 3, 4, and 5 years, using PFI as the primary endpoint based on the harmonized outcome definitions provided by the TCGA Pan-Cancer Clinical Data Resource (**Fig. 1A**)^25^.

**Fig. 1.**
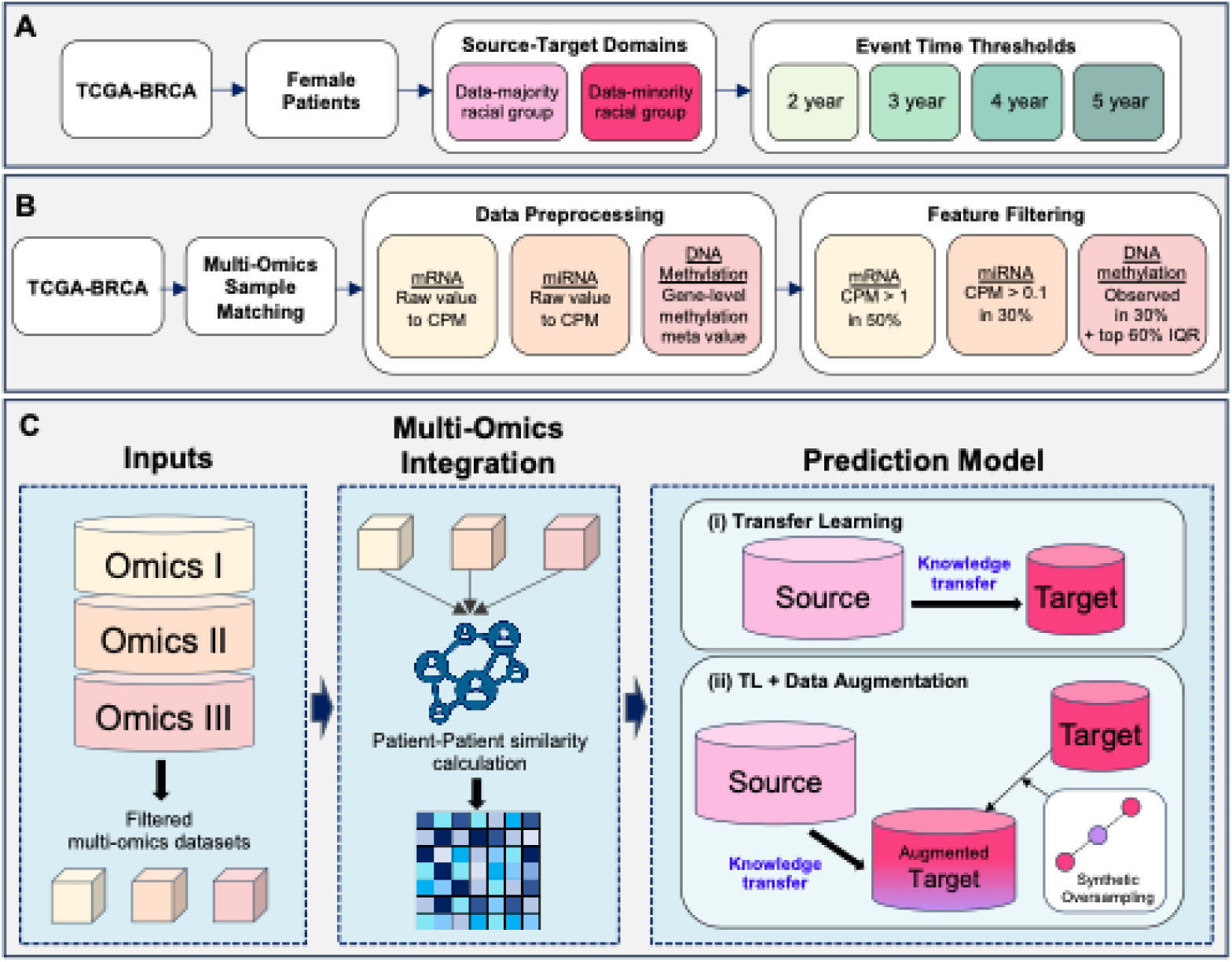
Overview of the MOTLAB framework for breast cancer health disparity reduction. Overview of the MOTLAB workflow from cohort definition and multi-omics processing to EA-to-AA transfer learning and PFI prediction. (**A**) Female TCGA-BRCA samples filtered to European American (EA) and African American (AA) groups for 2-, 3-, 4-, and 5-year PFI prediction. (**B**) mRNA expression, miRNA expression, and DNA methylation data were paired across shared samples, processed separately, and subjected to modality-specific feature filtering. (**C**) Filtered omics data were converted into patient-patient similarity representations and integrated across modalities. Conventional EA-to-AA TL and MOTLAB with SMOTE-based target-domain data augmentation were evaluated.

In this study, MOTLAB incorporated three molecular modalities—mRNA expression, miRNA expression, and DNA methylation—which represent transcriptional, post-transcriptional, and epigenetic information in BC (**Fig. 1B**)^26,27,28,29^. Following modality-specific preprocessing, each omics layer was represented by patient–patient similarities and integrated using optimized omics-specific weights to generate an integrated patient-level representation (**Fig. 1C**)^21,30^. Using this representation, we evaluated conventional CCSA-based TL from the EA source domain to the AA target domain and the final MOTLAB configuration, which additionally incorporated Synthetic Minority Over-sampling Technique (SMOTE)-based DA of the AA target domain data^24,31^. Following model development, we conducted a structured downstream to evaluate predictive performance, reliability, and biological relevance (**Fig. S1**). Thus, the key design of MOTLAB was to combine optimized multi-omics weighting, TL, and DA within a unified framework for PFI prediction in the data-minority group.

### 2.2 MOTLAB outperformed conventional learning strategies through weighted multi-omics integration and data augmentation

To evaluate the effectiveness of MOTLAB for BC PFI prediction, we compared four learning strategies: mixture learning trained on pooled samples and evaluated in AA, AA-specific independent learning, conventional TL, and MOTLAB. Model performance was evaluated using area under the receiver operating characteristic curve (AUROC) and balanced accuracy (BACC) across four PFI time settings. Across most settings, three-omics integration outperformed two-omics combinations (**Fig. 2**). At the 2-year PFI prediction, MOTLAB achieved an AUROC of approximately 0.70 and a BACC of approximately 0.61 with three-omics integration, compared with approximately 0.45–0.68 and 0.49–0.58, respectively, for the other models (**Fig. 2A, B**).

**Fig. 2.**
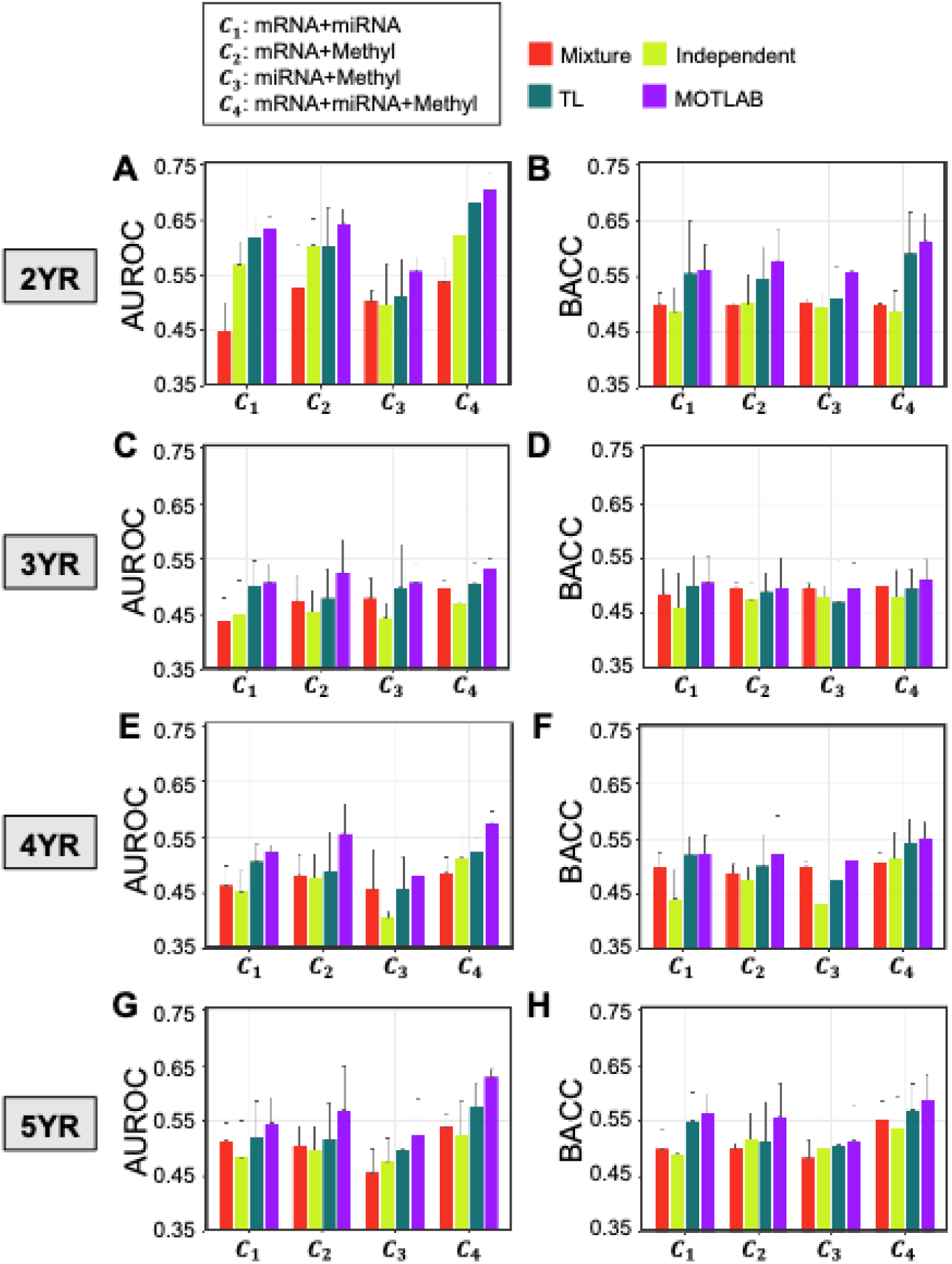
MOTLAB performance across multi-omics combinations and PFI predictions. AUROC and BACC for 2-(**A, B**), 3-(**C, D**), 4-(**E, F**), and 5-year (**G, H**) PFI prediction across four learning strategies. Omics combinations were C1, mRNA+miRNA; C2, mRNA+DNA methylation; C3, miRNA+DNA methylation; and C4, mRNA+miRNA+DNA methylation. Bars indicate mean performance across seed-level evaluations; error bars indicate SD.

Performance differences were more modest at 3 years, with AUROCs mostly ranging from approximately 0.44 to 0.53 and BACC values from approximately 0.46 to 0.50 (**Fig. 2C, D**). At 4 and 5 years, MOTLAB maintained stronger three-omics performance, achieving AUROCs of approximately 0.57 and 0.63 and BACC values of approximately 0.55 and 0.58, respectively (**Fig. 2E–H**). Comparisons across single-, two-, and three-omics settings showed that expanded omics data integration was associated with stronger MOTLAB performance (**Fig. S2**).

To assess the contribution of omics-specific weighting, we performed a grid search of weight combinations for individual omics data in multi-omics integration (**Fig. S3**). At the 2-year PFI prediction, ΔAUROC ranged from approximately −3.0% to +3.5%, with most weight combinations yielding positive values, while ΔBACC generally ranged from approximately −3.9% to +3.6% across PFI time settings (**Fig. S3A–B**). Across most weight combinations, MOTLAB outperformed conventional TL at the 3-, 4-, and, 5-year PFI predictions, with ΔAUROC ranging from-3.0% to +5.9%,-1.7% to +6.1%, and-1.9% to +5.6%, respectively, and corresponding ΔBACC ranges of-5.3% to +3.7% (**Fig. S3C-D**),-2.6% to +4.5% (**Fig. S3E-F**), and-3.1% to +4.4% (**Fig. S3G-H**). These results indicate that MOTLAB prediction performance was influenced by the breadth of multi-omics integration and the relative weighting of individual omics data.

### 2.3 MOTLAB enhanced predictive discrimination and calibration for PFI prediction

To benchmark MOTLAB against survival-model and TL baselines, we compared Cox elastic net (CoxEN), random survival forest (RSF), TL, and MOTLAB across all omics combinations and PFI time settings (**Fig. 3**)^32,33^. At the 2-year PFI prediction, MOTLAB achieved the highest AUROC across most multi-omics combinations, reaching approximately 0.70 with the three-omics combination compared with approximately 0.55 and 0.56 for CoxEN and RSF, respectively (**Fig. 3A**). BACC showed a similar pattern, with MOTLAB reaching approximately 0.62 compared with approximately 0.50–0.59 for the other models (**Fig. 3B**). Performance differences were more modest at 3 years, whereas three-omics MOTLAB maintained stronger performance at 4 and 5 years, with AUROCs of approximately 0.58 and 0.63 and BACC values of approximately 0.55 and 0.59, respectively (**Fig. 3C–H**). Patient-level paired bootstrap estimates of ΔAUROC generally favored MOTLAB over TL, particularly in several methylation-containing and three-omics settings (**Fig. S4**).

**Fig. 3.**
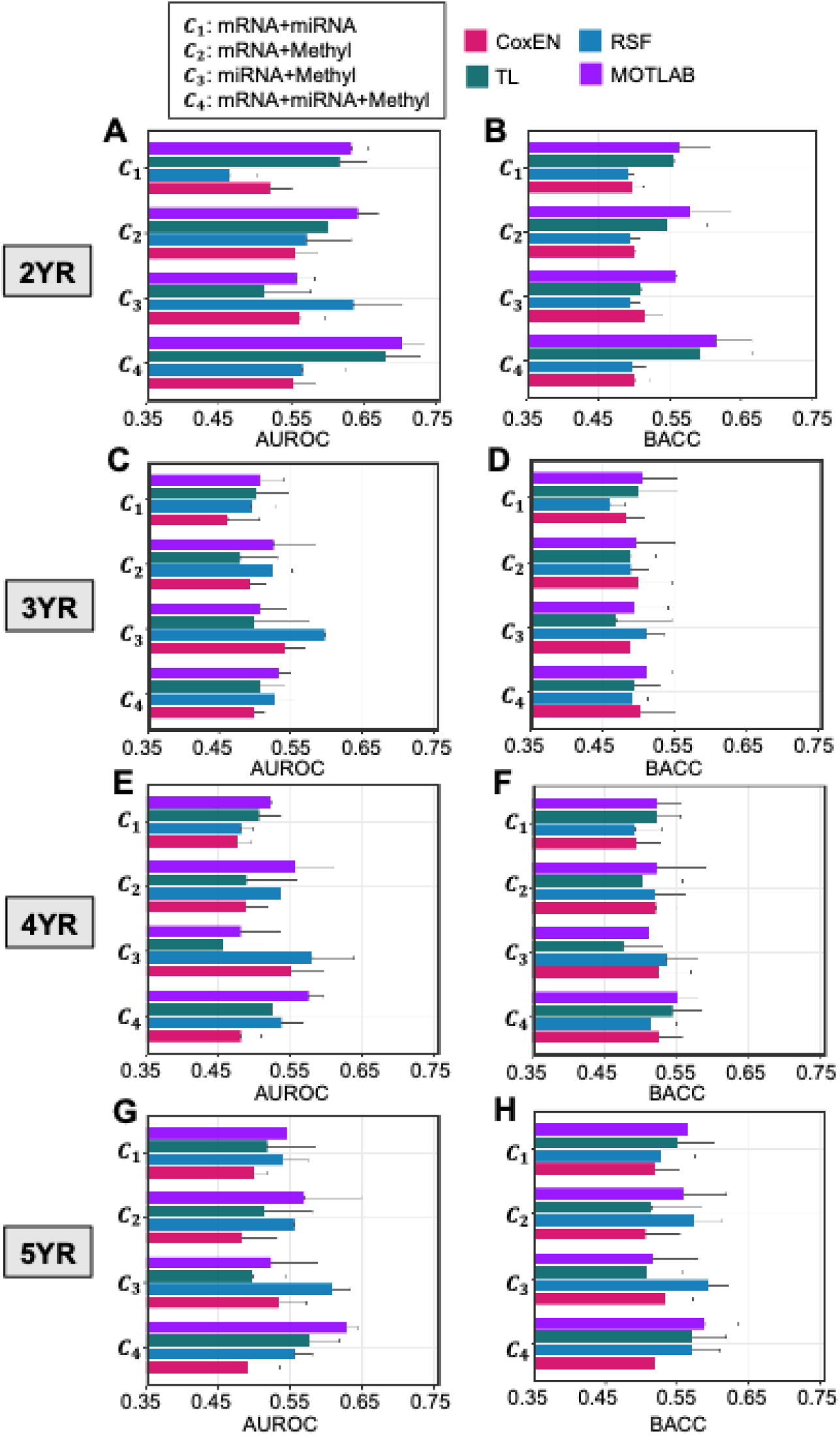
Benchmarking of MOTLAB against survival-model and transfer-learning baselines. AUROC and BACC for Cox elastic net (CoxEN), random survival forest (RSF), TL, MOTLAB, and comparator models across 2-(**A, B**), 3-(**C, D**), 4-(**E, F**), and 5-year (**G, H**) PFI prediction. Omics combinations were defined as in Fig. 2. Bars indicate mean performance across repeated evaluations; error bars indicate SD.

To complement AUROC and BACC, we evaluated discrimination, prediction error, and calibration using C-index, Brier score, and expected calibration error (ECE), respectively, across PFI time settings and omics combinations (**Fig. 4**)^34,35,36^. MOTLAB generally achieved the highest or near-highest C-index and lowest Brier scores than TL alone in three-omics integration across the all PFI predictions (**Fig. 4A–E**), with C-index and Brier score of approximately 0.64 and 0.18 at 2-year PFI predictions, respectively. ΔECE was predominantly negative at all PFI time settings (**Fig. 4F–I**). This improvement was strongest and most consistent at 2 years with all negative ΔECE values from −0.48 to −0.18 (**Fig. 4F**), and remained predominant at 3 years despite a few small positive values from −0.37 to +0.04 (**Fig. 4G**). At 4 and 5 years, ΔECE values remained from −0.18 to +0.01 and from −0.15 to +0.01, respectively (**Fig. 4H–I**). This result indicates lower calibration error for MOTLAB across most repetitions and omics combinations. Overall, MOTLAB performance is improved across discrimination, prediction error, and calibration relative to TL alone.

**Fig. 4.**
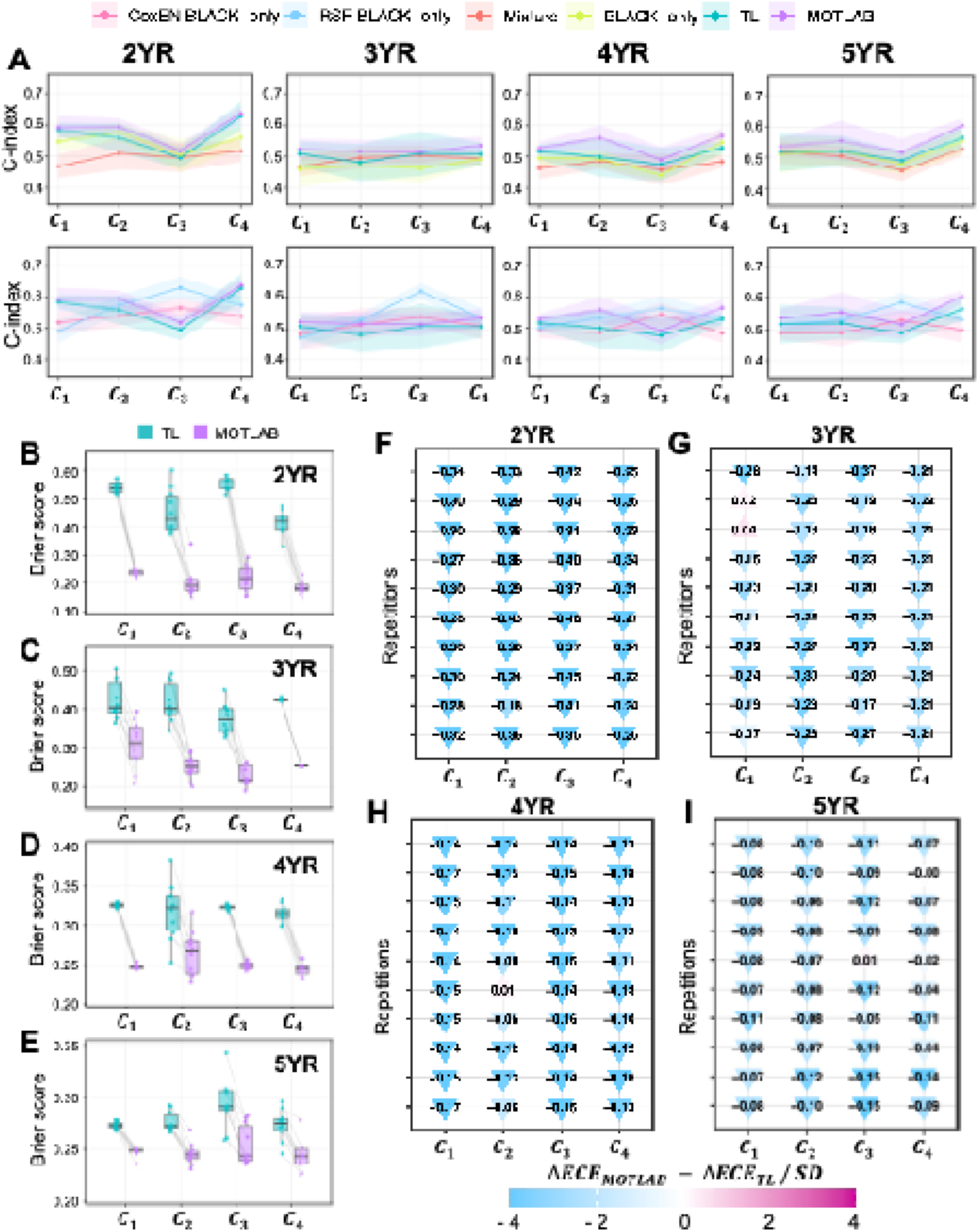
MOTLAB discrimination, prediction error, and calibration across PFI predictions. (**A**) C-index across 2-, 3-, 4-, and 5-year PFI predictions. The upper row compares CoxEN AA-only, RSF AA-only, TL, and MOTLAB; the lower row compares mixture, AA-only, TL, and MOTLAB models. Omics combinations are defined as in Fig. 3. Shaded regions indicate mean ± SD across seed-level evaluations. (**B–E**) Brier score distributions for TL and MOTLAB at 2 (**B**), 3 (**C**), 4 (**D**), and 5 years (**E**). Points indicate seed-level evaluations, boxplots summarize distributions, and gray lines connect paired evaluations from the same fold-derived seed. Lower values indicate lower prediction error. (**F–I**) ΔECE between MOTLAB and TL at 2 (**F**), 3 (**G**), 4 (**H**), and 5 years (**I**), defined as ECE_MOTLAB − ECE_TL. Triangle fill represents ΔECE divided by the within-year SD without mean-centering; labels show unscaled ΔECE. Negative values indicate lower calibration error for MOTLAB.

### 2.4 MOTLAB maintained robust PFI prediction and risk stratification across BC molecular and clinical subtypes

To evaluate robustness of MOTLAB in molecular and clinical subtypes, we incorporated molecular and clinical annotation data, including PAM50 intrinsic subtype, tumor stage, tumor size, lymph node status, and metastasis status^26,37^. Because calculation of performance estimates is influenced by sample number of individual subgroups, we summarized event and non-event composition within individual subgroups across PFI time settings (**Fig. S5**). We evaluated subgroup performance differences between MOTLAB and TL alone using ΔAUROC, ΔBACC, ΔC-index, and ΔBrier score in individual subgroups (**Fig. 5**). Across PAM50 subtypes, HER2-enriched tumors showed ΔAUROC values of approximately 0.09, 0.11, 0.06, and 0.08 across 2-, 3-, 4-, and 5-year PFI settings, respectively (**Fig. 5A**). Brier score improvements were particularly strong across PAM50 subtypes, with negative ΔBrier values observed in most subtypes and PFI time settings (**Fig. 5A**). In case of tumor stage, stage IV tumors showing a ΔAUROC of approximately 0.07 at 2-year PFI and stage III and stage II tumors showing ΔAUROC values of approximately 0.04 and 0.03, respectively (**Fig. 5B**). Tumor size, lymph node, and metastasis subgroups also showed that MOTLAB maintained improved performance across most subgroups and PFI time settings (**Fig. 5C–E**).

**Fig. 5.**
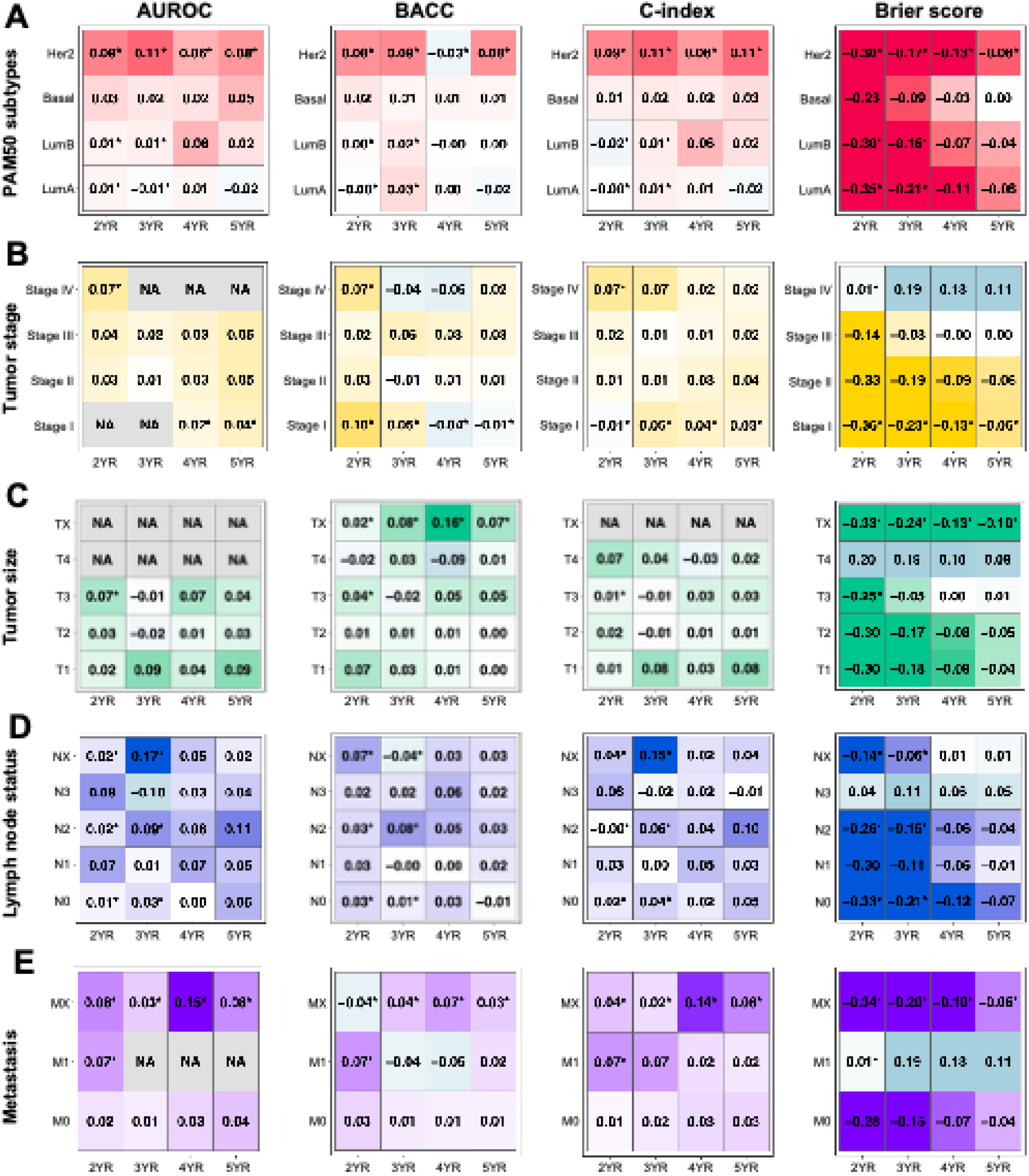
MOTLAB performance across molecular and clinical subgroups. Heatmaps show subgroup-specific differences between MOTLAB and TL across 2-, 3-, 4-, and 5-year PFI predictions for PAM50 subtype (**A**), stage (**B**), tumor size (**C**), lymph node status (**D**), and metastasis status (**E**). Δmetrics were calculated as MOTLAB − TL. Positive ΔAUROC, ΔBACC, and ΔC-index and negative ΔBrier score favor MOTLAB. NA indicates unavailable or non-evaluable paired estimates. Asterisks indicate subgroup-PFI combinations with mean event count <3.

To examine whether MOTLAB-derived risk groups were clinically interpretable, we compared PFI event composition and restricted mean survival time (RMST) differences between MOTLAB low-risk and high-risk groups (**Fig. 6**). RMST estimates the expected event-free survival time up to a specified PFI time and provides an absolute time-scale summary that can be easier to interpret clinically than a hazard ratio, particularly when proportional hazards assumptions may be questionable^38,39^. At the 2-year PFI prediction, the low-risk group included 51 samples, and the high-risk group included 52 samples, with PFI events less frequent in the low-risk group than in the high-risk group, approximately 14% versus 37%, respectively (**Fig. 6A**). Similar patterns were observed at 3-, 4-, and 5-year predictions, with event rates of approximately 26% versus 36%, 28% versus 44%, and 33% versus 53% in low-risk versus high-risk groups, respectively (**Fig. 6C, E, G**). RMST analyses supported the robustness of MOTLAB-derived risk stratification across molecular and clinical subgroups. MOTLAB low-risk patients generally had longer mean PFI than predicted high-risk patients, with positive RMST differences observed across most PAM50, stage, tumor, lymph node, and metastatic subgroups (**Fig. 6B, D, F, H**). The overall RMST difference remained positive across all PFI time settings, and the direction of separation was maintained in many individual subgroups despite variability in effect size and confidence intervals. These results indicate that MOTLAB-derived risk groups captured risk differences across PFI time settings beyond the overall cohort and across heterogeneous subtypes. Overall, these findings support the potential clinical utility of MOTLAB for risk stratification in heterogeneous BC patients.

**Fig. 6.**
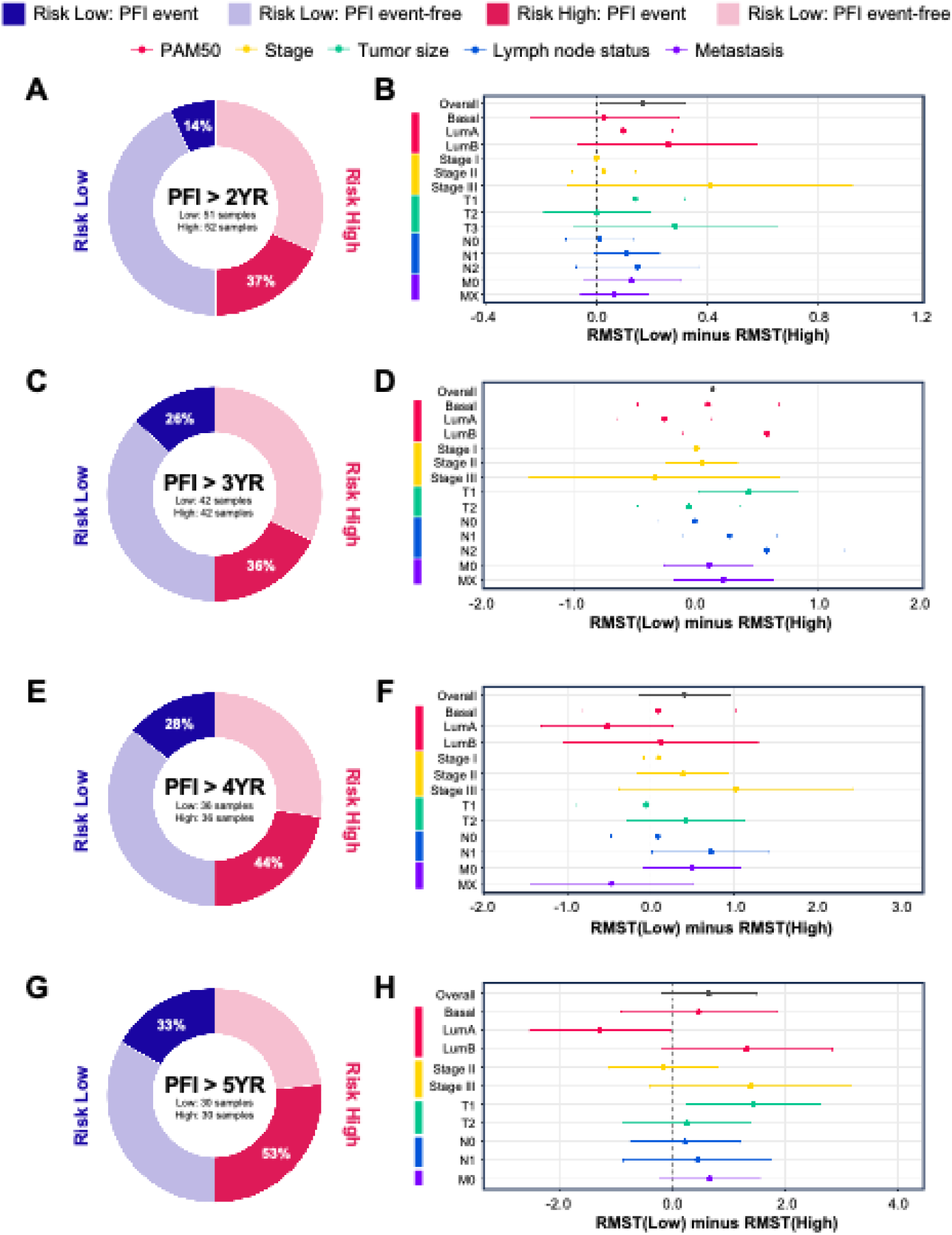
MOTLAB-derived risk groups differed in PFI event burden and restricted mean survival time. MOTLAB low-and high-risk groups were evaluated at 2 (**A, B**), 3 (**C, D**), 4 (**E, F**), and 5 years (**G, H**). Donut charts show PFI event and non-event composition; centers indicate sample size. Risk groups were defined by the median MOTLAB risk score at each PFI time setting. Forest plots show RMST differences (RMST_low − RMST_high) for the overall cohort and eligible subgroups. Positive values indicate longer restricted mean PFI in the low-risk group. Points indicate RMST differences; horizontal lines indicate 95% confidence intervals.

### 2.5 MOTLAB risk scores captured breast cancer-relevant molecular programs across multi-omics data

To investigate whether MOTLAB-derived risk scores capture biologically interpretable signals, we calculated correlations between risk scores and molecular features, including MSigDB Hallmark pathway score, miRNA abundance, gene-level methylation, across PFI time settings (**Fig. 7**). This analysis was designed to determine whether MOTLAB-derived risk scores in AA patients were associated with BC-related molecular pathways. We used the Hallmark gene set collection from MSigDB because it summarizes redundant gene sets into coherent biological states and processes such as proliferation, immune response, DNA damage, metabolism, and oncogenic signaling^40^. In parallel, we annotated miRNAs using the Human MicroRNA Disease Database (HMDD v4.0), a curated resource of experimentally supported associations between human miRNAs and diseases, to identify miRNAs linked to BC^41^.

**Fig. 7.**
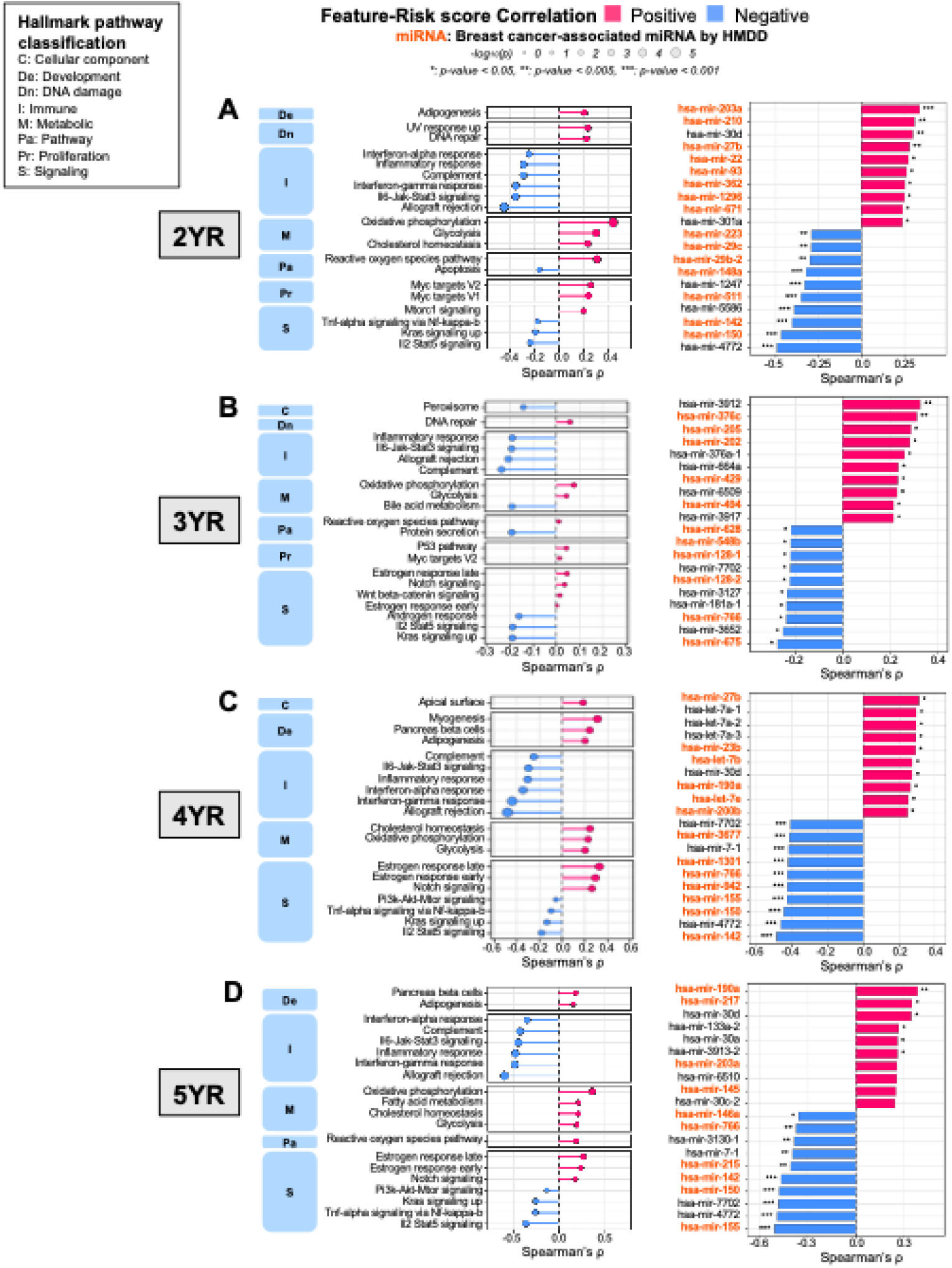
MOTLAB risk scores were associated with Hallmark pathway and miRNA programs. Spearman correlations between MOTLAB risk scores and Hallmark pathway scores or miRNA abundance at 2 (A), 3 (B), 4 (C), and 5 years (D). Left panels show Hallmark pathways grouped by process category; right panels show risk-associated miRNAs. Red and blue indicate positive and negative correlations with MOTLAB risk, respectively. Asterisks indicate nominal Spearman p values (*p < 0.05, **p < 0.005, ***p < 0.001). Orange labels indicate BC-associated miRNAs identified from HMDD v4 disease entries containing “breast.”

At the 2-year PFI prediction, higher MOTLAB risk was associated with several Hallmark pathways linked to aggressive BC, including mTORC1 signaling^42^, MYC targets V1/V2^43^, reactive oxygen species (ROS)^44^, glycolysis^45^, and oxidative phosphorylation^46^ (**Fig. 7A**). BC studies have linked increased MYC target activity^43^, ROS abundance^44^, and oxidative phosphorylation-associated gene expression^46^ to aggressive BC or poor survival in BC, while mTORC1 signaling^42^ and glycolysis-related signatures^45^ have also shown subtype-specific prognostic associations. These findings support the biological relevance of the molecular variation captured by MOTLAB risk. In contrast, lower risk score was associated with interferon-and immune-related pathways, including interferon-α response, interferon-γ response, inflammatory response, complement, and allograft rejection (**Fig. 7A**). This pattern is consistent with subtype-specific evidence linking type I interferon activity to favorable outcomes^47^, IFN-γ signaling to prognosis^48^, and enhanced tumor immune activity to improved survival in TNBC and HER2-positive BC^49,50^. This pattern is consistent with evidence linking greater antitumor immune activity and interferon-related transcriptional pathways to better outcomes in BC. Regarding miRNA abundance, 2-year MOTLAB-predicted risk showed positively correlated HMDD-annotated miRNAs including hsa-miR-203a, hsa-miR-210, hsa-miR-27b, hsa-miR-93, hsa-miR-362, and hsa-miR-22, with positive correlations reaching approximately 0.3–0.4 (**Fig. 7A**)^41,51,52,53,54,55,56^. Conversely, higher MOTLAB risk was negatively correlated with several HMDD-annotated miRNAs, including miR-29c, miR-148a, miR-142, and miR-150, whose higher expression has been associated with tumor-suppressive activity or more better outcomes in previous BC studies^57,58,59,60^. These results suggest that MOTLAB-derived risk scores capture

BC-relevant post-transcriptional regulatory variation. Across the 3-, 4-, and 5-year PFI predictions, molecular associations recapitulated the pattern observed at 2-year PFI prediction, with higher MOTLAB risk primarily associated with metabolic and tumor-related programs and lower MOTLAB risk associated with immune-related pathways (**Fig. 7B–D**). Oxidative phosphorylation and glycolysis correlated with risk scores from all three PFI predictions, while estrogen-response and Notch signaling became more prominent at 4 and 5 years. Conversely, inflammatory, complement, allograft-rejection, and interferon-related programs showed recurrent inverse associations with MOTLAB risk. Consistent with findings from pathway analysis, HMDD-annotated BC-associated miRNAs continued to appear among the strongest positive and negative risk correlations at these PFI time settings. We additionally investigated entire cohort heatmaps across all PFI time settings, which showed that risk-score-associated Hallmark pathways and miRNAs reflected structured molecular patterns across MOTLAB-defined risk groups and clinical annotations (**Fig. S6**). The ranked methylation-correlation profiles showed both positively and negatively risk-associated gene-level methylation patterns (**Fig. S7**). Overall, these findings suggest that MOTLAB-derived risk scores that remained informative across molecular and clinical subgroups and were associated with BC-relevant pathway, miRNA, and DNA methylation patterns, supporting their clinical and biological interpretability.

## Discussion

In this study, we developed MOTLAB, a weighted multi-omics TL framework designed to improve BC outcome prediction under multi-omics data from data-minority ancestry group. By optimizing the contributions of individual omics data, transferring knowledge from the EA source domain to the AA target domain, and augmenting the AA data, MOTLAB was developed to address limitations in molecular representation for reducing BC disparities. Across four PFI prediction tasks, MOTLAB generally achieved better performance than conventional ML approaches and survival model benchmarks, while showing lower prediction error and improved calibration relative to TL alone. Moreover, MOTLAB showed the robustness of performance across molecular and clinical subgroups, and MOTLAB-derived risk scores stratified AA patients by PFI and were associated with BC-relevant molecular features. Overall, MOTLAB made methodological advance in how these ML strategies are integrated rather than in any individual component. Previous works established TL as a strategy for leveraging information from data-majority groups to improve prediction in data-minority groups^14,61^. Patient-similarity approaches have demonstrated the utility of integrating heterogeneous molecular data whereas SMOTE was originally developed to overcome class imbalance^21,30,31^. MOTLAB extends these concepts by optimizing the contribution of each omics modality and by incorporating DA as an additional approach to overcome limited representation of data-minority group. The best performance observed with multi-omics integration and optimized weighting is consistent with prior evidence that complementary information across multi-omics data can improve cancer outcome modeling^26,62,63^.

The molecular analyses support the biological relevance of the risk information learned by MOTLAB. MOTLAB-derived risk scores were associated with established MSigDB Hallmark pathways, breast cancer-associated miRNAs curated in HMDD, and gene-level DNA methylation patterns, linking model-derived risk to transcriptional, post-transcriptional, and epigenetic variation relevant to breast cancer biology^26,40,41^. This multi-omics context is relevant given previous evidence of molecular heterogeneity in BC and ancestry-associated variation across transcriptomic and epigenetic features in TCGA cohorts^64,65^. However, previous ancestry-based molecular analyses have also highlighted the importance of tumor subtype and technical factors when interpreting such differences^65^. Accordingly, these findings provide biological context for the predictive signals and support the interpretation of the identified pathways, miRNAs, and methylation features as molecular correlates of PFI. Beyond discrimination, the lower prediction error and improved calibration observed relative to TL alone provide additional information about model performance. Discrimination quantifies the ability to distinguish patients with different outcomes, whereas calibration evaluates agreement between predicted and observed risks; both are important dimensions of predictive model evaluation^66,67^. The consistently improved predictive performance across molecular and clinical subgroups supports the robust applicability of MOTLAB across diverse BC characteristics. With these findings, MOTLAB provide a generalizable framework for improving personalized prediction in BC data-minority groups.

To improve the robustness and effectiveness of MOTLAB, several additional considerations should be addressed in future studies aimed at reducing BC disparities. The analysis was based on a single TCGA cohort with a relatively small racial group and therefore requires validation in independent, racially diverse multi-omics cohorts. Moreover, the paired-bootstrap confidence intervals between MOTLAB and TL alone AUROC differences included zero, which is indicating that the observed performance advantages should be interpreted as directional rather than definitive evidence of statistical superiority. The time-specific binary PFI prediction also simplifies the complete censored time-to-event. Future work should therefore evaluate MOTLAB in larger multi-institutional cohorts, extend the framework to native survival-learning objectives, and integrate molecular, clinical, and treatment features.

## Methods

### TCGA-BRCA cohort and ancestry annotation

We analyzed patients with breast invasive carcinoma from The Cancer Genome Atlas. Multi-omics and clinical outcome data were obtained from TCGA resources through the National Cancer Institute Genomic Data Commons, which provides harmonized cancer genomic and clinical data infrastructure for TCGA and related programs^68^. Genetic ancestry labels were obtained from TCGA ancestry resources that infer ancestry from germline genetic variation^65,69^. Patients annotated as EA were mapped to the EA group, and patients annotated as AA were mapped to the AA group. The present study focused on these two ancestry groups because the modeling objective was to evaluate TL from a data-majority group to a data-minority group in BC prognosis. Throughout the modeling analyses, the AA group refers to patients assigned to AA based on germline genetic ancestry annotation. We therefore interpret the analysis as addressing a racial-disparities-related data-minority prediction setting, while recognizing that genetic ancestry and socially defined race are related but non-identical constructs.

### Clinical endpoint

PFI was used as the primary clinical endpoint and was obtained from the TCGA Pan-Cancer Clinical Data Resource, which harmonized multiple survival endpoints across TCGA cancer types and proposed PFI as a clinically practical endpoint when disease progression or recurrence is more informative than death alone^25^. For each experiment, PFI time and event indicators were converted into binary labels at 2-, 3-, 4-, or 5-year predictions.

### Cohort inclusion and exclusion criteria

Patients were included if they had available mRNA expression, miRNA expression, DNA methylation, genetic ancestry annotation, and PFI outcome data. TCGA barcodes were harmonized at the patient by truncating sample identifiers to the first 12 characters, and duplicate patient entries were removed by retaining the first occurrence. For three-omics analyses, only patients shared across all three omics matrices were retained. Patients were excluded from a time-specific analysis if they were censored on or before the time setting, because their time-specific event status could not be assigned unambiguously.

### Modeling cohort construction and endpoint definition

After ancestry filtering and omics matching, the analytic cohort was restricted to EA and AA patients only. Survival labels were defined for PFI prediction of 2-, 3-, 4-, or 5-years. For a given PFI time *t*, patients censored on or before *t* were excluded from modeling. Patients with an event by *t* were labeled as 0, whereas those remaining event-free beyond t were labeled as 1.

This time-specific formulation was used throughout the TL and baseline pipelines and was paired with supplementary discrimination, calibration, and risk-ranking analyses to mitigate the information loss inherent in binary discretization of survival outcomes.

### Multi-omics integration

The proposed framework integrated mRNA, miRNA, and DNA methylation using a weighted multi-omics representation. Let *x*^(*mRNA*)^, *x*^(*miRNA*)^, and *x*^*(methyl)*^ denote the sample-by-feature matrices for the three omics layers. Within each omics matrix, values were converted to numeric format, missing values were imputed using the sample-wise mean, and each sample vector was normalized by subtracting its own mean and dividing by its own standard deviation. This preprocessing was performed separately within each omics layer.

The integrated representation was then constructed by weighted concatenation:

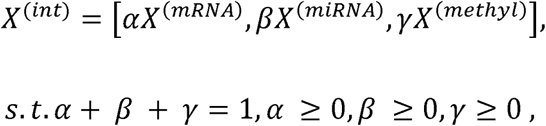

Thus, the weights controlled the relative contribution of each omics modality. In the three-omics experiments, candidate weight combinations were generated on a discrete grid with step size 0.1 and minimum weight 0.1, yielding combinations satisfying the simplex constraint.

In the TL pipeline, the weighted integrated matrix was transformed into an anchor-based Pearson correlation coefficient (PCC) representation within each cross-validation split. For each query sample, PCC values were computed against an anchor set consisting of all source domain EA samples and the few-shot target domain AA training samples. Therefore, the final TL input was an anchor-similarity representation derived from the weighted integrated omics profile.

### Transfer learning framework

We defined EA patients as the source domain and AA patients as the target domain. The TL framework was implemented using Contrastive Classification Semantic Alignment (CCSA), a supervised domain adaptation strategy that aligns samples from different domains in a shared latent space while preserving class-discriminative structure^24^. The model consisted of a shared encoder applied to paired source and target inputs, followed by a softmax classification head and a contrastive semantic alignment branch. The encoder was instantiated through *Initialization_kernel.Create_Model(hiddenLayers=[100], dr=dr)*, and encoder weights were shared across the two Siamese branches. A two-node softmax classifier generated the probability of the binary prognosis label, while a contrastive distance term was computed between paired latent representations. The model was trained by minimizing a weighted sum of classification loss and CCSA loss:

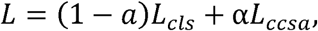

where L_*cls*_ is the sparse categorical cross-entropy loss for prognosis classification and L_*ccsa*_ is the contrastive semantic alignment loss for domain adaptation^24^. The domain adaptation loss weight was fixed at a = 0.1. The TL procedure should therefore be understood as joint supervised domain adaptation with few labeled target samples.

### Few-shot target sampling

Model evaluation used repeated 3-fold stratified cross-validation with 10 random seeds in the three-omics TL experiments. For each seed, AA patients were partitioned into three stratified folds according to the binary outcome label. Within each training fold, up to five AA samples from each outcome class were selected as the few-shot target training subset. The remaining AA samples from the training fold were assigned to the validation subset, and the held-out fold served as the AA test subset. The anchor set used for PCC-based transfer features included only

EA source samples and target-train AA samples. Target-validation and target-test samples were not included in the anchor set, preventing information leakage from validation or test samples into feature construction.

### Feature selection and optimization

Within each fold, after construction of the anchor-based PCC representation, univariate feature selection was performed using *SelectKBest* with the ANOVA F-statistic from *scikit-learn*.

Feature selection was fit using source domain EA training labels only, and the selected features were applied to source, target-train, target-validation, and target-test data within the same fold. In the reported three-omics experiments, the number of selected features was searched across {50, 100, 150, 200, 300, 500}. The model was trained using stochastic gradient descent with Nesterov momentum. The learning rate was 0.001, momentum was 0.9, dropout was 0.0 for the shown three-omics runs, batch size was 10, learning-rate decay was 0, and early stopping patience was 10 epochs. Model selection was performed using the validation subset, and the final event-class threshold was tuned on validation data by maximizing the F1 score before application to the independent AA test subset.

### Data augmentation using Synthetic Minority Over-sampling Technique

To improve learning stability in the target domain, we implemented a TL with DA variant (TLDA), in which only the target-train AA subset was resampled before CCSA training. Oversampling was performed after construction of the anchor-based transfer representation and feature selection, but before pair generation and model fitting. Validation and test subsets were never oversampled. SMOTE was used when both target outcome classes were present and the minority class size was at least 2^31^. SMOTE parameters were determined dynamically within each fold by setting the desired target sample counts to approximately match the source sample size while preserving the observed target class proportion. The number of nearest neighbors was set adaptively as k_*SMOTE*_ = min(5, n_*BLACK*_ - 1) where n_*BLACK*_ denotes the number of target-train samples in the smaller class for that fold. When the minority class size was too small for SMOTE, random oversampling was performed. SMOTE operations were logged to fold-specific JSON files, including class counts before and after resampling, the selected k-neighbors value, and whether the resampled target size matched the source size.

### Conventional machine learning baselines

To contextualize the proposed TL approach, we implemented two non-transfer neural baselines for time-specific PFI prediction: mixture learning and AA-specific independent learning. In the mixture baseline, EA and AA patients were pooled during model training, and performance was evaluated in AA held-out test samples. In the AA-only independent baseline, models were trained and evaluated only within AA patients. The neural baseline model was a multilayer perceptron with one hidden layer of 100 units and a two-node softmax output. Class-weighted negative log-likelihood was used to address outcome imbalance. Scaling and feature selection were performed within each training fold.

### Cox elastic-net and random survival forest

Classical survival baselines were implemented to compare the proposed TL framework with established right-censored survival modeling approaches. Cox elastic-net regression (CoxEN) was used as a penalized Cox proportional hazards model suitable for high-dimensional omics predictors^32,70,71^. Random survival forest (RSF) was used as a non-parametric ensemble survival method for right-censored data^33^. For CoxEN and RSF, omics inputs were constructed for mRNA+miRNA, mRNA+methylation, miRNA+methylation, and mRNA+miRNA+methylation combinations. Each omics layer was row-normalized and combined with equal weights within the relevant combination. Within each fold, missing values were imputed using training-fold medians, features were standardized using training-fold means and standard deviations, and the top 300 features were selected using an ANOVA F statistic fit only in the training data. CoxEN and RSF were trained using survival time and event indicators from the training fold, and predictions for held-out AA patients included a survival risk score and time-specific event probability. CoxEN was fit using cross-validated penalized Cox regression with elastic-net mixing parameter 0.5. RSF was fit using 300 trees, square-root feature subsampling, and node size 5. Seed-level performance was computed from concatenated out-of-fold AA predictions.

### Performance evaluation

The primary analysis focused on predictive performance in the AA test cohort, because the main goal was to improve clinical outcome prediction for the data-minority group. Time-specific discrimination was evaluated using the area under the receiver operating characteristic curve (AUROC) and balanced accuracy (BACC), whereas time-to-event discrimination was assessed using Harrell’s concordance index (C-index) based on observed PFI times, event indicators, and predicted risk scores. Calibration was quantified using the Brier score, expected calibration error (ECE), maximum calibration error (MCE), and calibration-bin summaries. Subgroup analyses were conducted using PAM50 intrinsic subtype, tumor stage, tumor size, lymph node status, and metastasis status from the subgroup annotation table. Performance and calibration summaries were generated at both the overall and subgroups across repeated cross-validation seeds.

### Patient-level paired bootstrap analysis

To evaluate the robustness of the observed performance differences between MOTLAB and conventional TL, patient-level paired bootstrap resampling was performed for individual PFI time settings and omics combinations. Bootstrap analyses were restricted to MOTLAB and TL because patient-level prediction scores were available for both models. For each patient, prediction scores were first averaged across matched random seeds shared between the two models. Outcome-stratified paired bootstrap resampling was then performed by sampling patients with replacement while preserving the observed event and non-event composition. For each bootstrap replicate, AUROC was calculated separately for MOTLAB and TL, and the paired difference was summarized as ΔAUROC = AUROC_MOTLAB − AUROC_TL. A total of 10,000 bootstrap replicates were generated for each comparison. Bias-corrected and accelerated 95% confidence intervals were estimated using the boot package in R. Bootstrap summaries additionally included the proportion of replicates with ΔAUROC > 0 and Monte Carlo standard errors for these proportions.

### Risk score definition, risk stratification, and restricted mean survival time analysis

For downstream risk-stratification and biological interpretation analyses, MOTLAB-derived risk scores were calculated from model-predicted class probabilities. Because the binary label was encoded as 0 for patients with a PFI event by the PFI time setting and 1 for patients remaining event-free beyond the PFI time setting, the MOTLAB risk score was defined as one minus the predicted probability of class 1. For each PFI prediction, patients were divided into predicted low-risk and high-risk groups using the median MOTLAB risk score. PFI event and non-event composition was summarized within each risk group at the 2-, 3-, 4-, and 5-year predictions.

Restricted mean survival time (RMST) was calculated up to the corresponding prediction to provide an absolute time-scale summary of clinical separation between predicted risk groups. RMST differences were summarized as RMST_low-risk − RMST_high-risk, such that positive values indicated longer restricted mean PFI in the predicted low-risk group. RMST differences were also calculated within molecular and clinical subgroups when sufficient samples were available. Confidence intervals for RMST differences were estimated using the unadjusted RMST difference output from the survRM2 package in R, and subgroup estimates with limited sample size, or event counts were interpreted cautiously.

### Subgroup performance and outcome-composition analyses

Subgroup analyses were conducted using PAM50 intrinsic subtype, tumor stage, tumor size, lymph node status, and metastasis status from the clinical and molecular annotation table. For each subgroup, PFI time setting, and performance metric, MOTLAB was compared with TL alone using paired performance differences. ΔAUROC, ΔBACC, and ΔC-index were defined as the MOTLAB value minus the TL alone value, whereas ΔBrier score was defined analogously such that negative values indicated lower prediction error for MOTLAB. To contextualize subgroup performance estimates, event and non-event composition was summarized within each molecular and clinical subgroup across the 2-, 3-, 4-, and 5-year PFI predictions. Composition plots counted unique patients only and did not count repeated prediction rows across seeds, models, or omics-combination files. Subgroup-PFI time combinations with low mean event counts were flagged in the subgroup performance heatmaps and interpreted cautiously. Metrics that could not be estimated, for example because only one outcome class was present, were reported as not available.

### Calibration analysis and **Δ**ECE definition

Calibration was evaluated to determine whether predicted probabilities agreed with observed time-specific PFI event status. The Brier score was used as a proper scoring rule for prediction error, with lower values indicating more accurate probabilistic predictions. ECE was calculated by grouping predicted probabilities into 10 calibration bins and summing the bin-size-weighted absolute difference between the mean predicted risk and the observed event frequency within each bin. MCE was defined as the maximum absolute bin-level difference between predicted and observed risk across calibration bins. For paired calibration comparisons between MOTLAB and TL alone, ΔECE was defined as ECE_MOTLAB − ECE_TL. Therefore, negative ΔECE values indicated lower calibration error for MOTLAB. ΔECE values were calculated within each PFI prediction, omics-combination setting, and repeated cross-validation seed whenever paired MOTLAB and TL alone predictions were available.

### Multi-omics analysis of MOTLAB risk scores

To assess whether MOTLAB-derived risk scores were associated with biologically interpretable molecular programs, we correlated MOTLAB risk scores with Hallmark pathway activity, miRNA abundance, and gene-level DNA methylation summaries across four PFI predictions.

Hallmark gene sets were obtained from the MSigDB Hallmark collection^40^, and pathway scores were calculated from z-scored mRNA expression values for genes available in each gene set. miRNA features were annotated using HMDD v4.0 to identify experimentally supported BC-associated miRNAs^41^. For DNA methylation analysis, probe-level methylation features were summarized at the gene-level by averaging available methylation values mapped to each gene. Spearman correlations were calculated between MOTLAB risk scores and each Hallmark pathway score, miRNA feature, or gene-level methylation summary at each PFI prediction. P-values were calculated for Spearman correlations. Patient-level heatmaps and ranked correlation plots were used to visualize risk-associated pathway, miRNA, and methylation patterns. These analyses were interpreted as molecular context for MOTLAB-derived risk scores. Whole-cohort heatmaps were generated by ordering patients according to the MOTLAB-derived risk score within each PFI time setting. Hallmark pathway scores and HMDD-annotated miRNA abundances were displayed using row-wise standardized values to visualize global molecular patterns across predicted risk groups.

## Data availability

The multi-omics and clinical outcome data used in this study were derived from The Cancer Genome Atlas breast invasive carcinoma cohort. TCGA molecular and clinical resources are available through the National Cancer Institute Genomic Data Commons (https://portal.gdc.cancer.gov/). MOTLAB is available at https://github.com/wan-lab/MOTLAB.

## Acknowledgements

Research reported in this publication was supported by the U.S. National Science Foundation under Award Numbers 2500836 and 2614824, the National Cancer Institute of the National Institutes of Health under Award Number R03CA317707, the National Institute Of Alcohol Abuse And Alcoholism of the National Institutes of Health under Award Number R21AA032098, and the Office Of The Director, National Institutes Of Health of the National Institutes of Health under Award Number R03OD038391. This research was supported by the State of Nebraska through the Pediatric Cancer Research Group, part of the Child Health Research Institute. This work was also partially supported by the University of Nebraska Collaboration Initiative Grant from the Nebraska Research Initiative (NRI). The content is solely the responsibility of the authors and does not necessarily represent the official views of the funding organizations.

## Author contributions

M.B.: Data collecting and preprocessing, model development and optimization, data analysis and interpretation, manuscript preparation, editing and review. L.L.: Data collecting and preprocessing, script validation. V.B.: Manuscript review. J.W.: Manuscript editing and review. S.W.: Study conceptualization, design, and supervision, manuscript editing and review.

## Declaration of interests

The authors declare no competing interests.

## Notes

### Competing Interest Statement

The authors have declared no competing interest.

### Summary of Updates

All figures, supplemental files, result sections updated.

