## Supplemental Figures for "MOTLAB: A Weighted Multi-Omics Transfer Learning Framework for Reducing Racial Disparities in Breast Cancer"

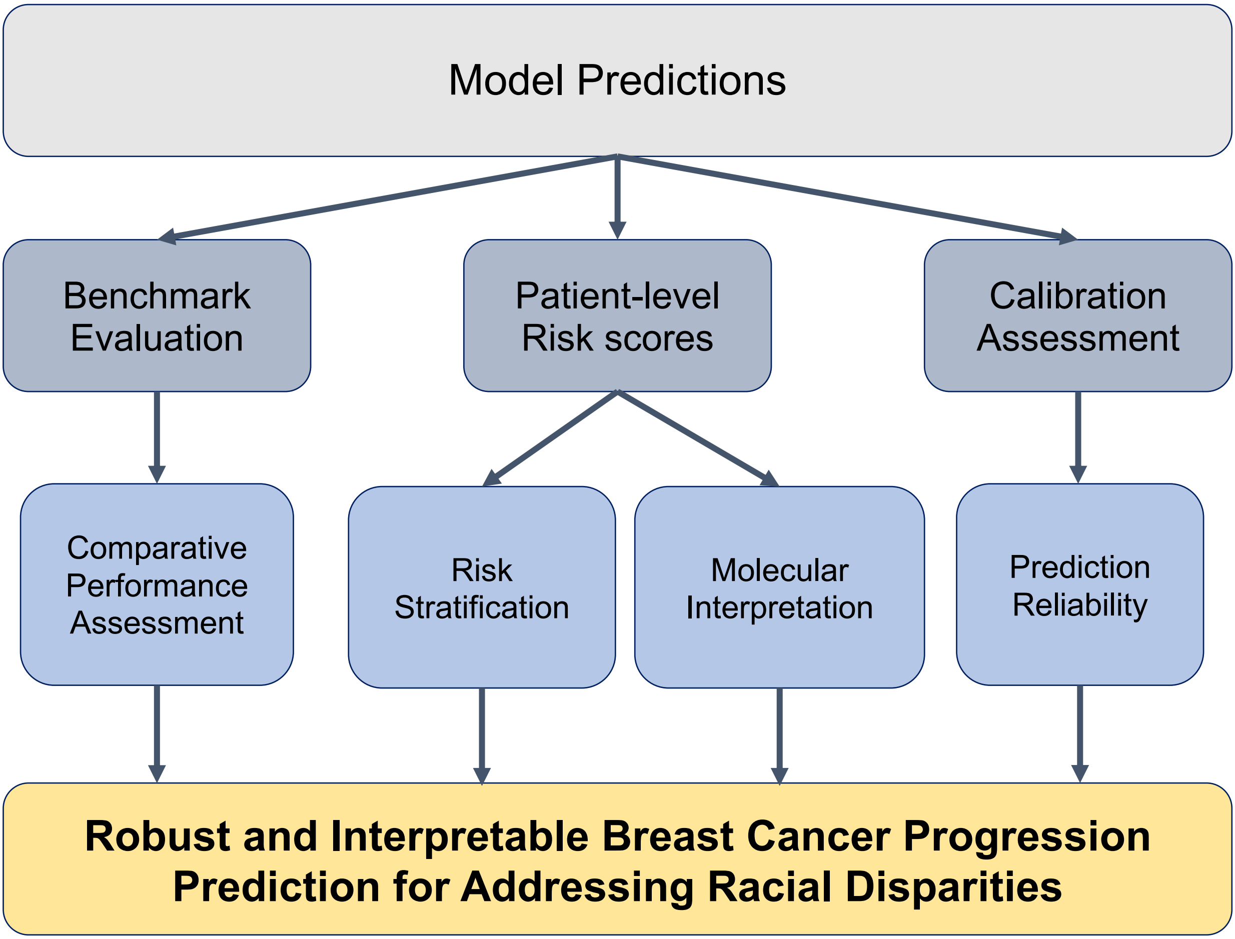


**Figure S1. Workflow of downstream analyses for comprehensive evaluation of MOTLAB predictions.**
Following model prediction, benchmark evaluation was performed to compare predictive performance against baseline models. MOTLAB-derived risk scores were subsequently used for clinical risk stratification and molecular interpretation on patient-level. Calibration assessment was conducted to evaluate the agreement between predicted and observed event probabilities and to assess prediction reliability. Thus, these downstream analyses provide a comprehensive evaluation of predictive performance, calibration, clinical utility, and biological interpretability of MOTLAB in the data-minority group.

**Figure S2**

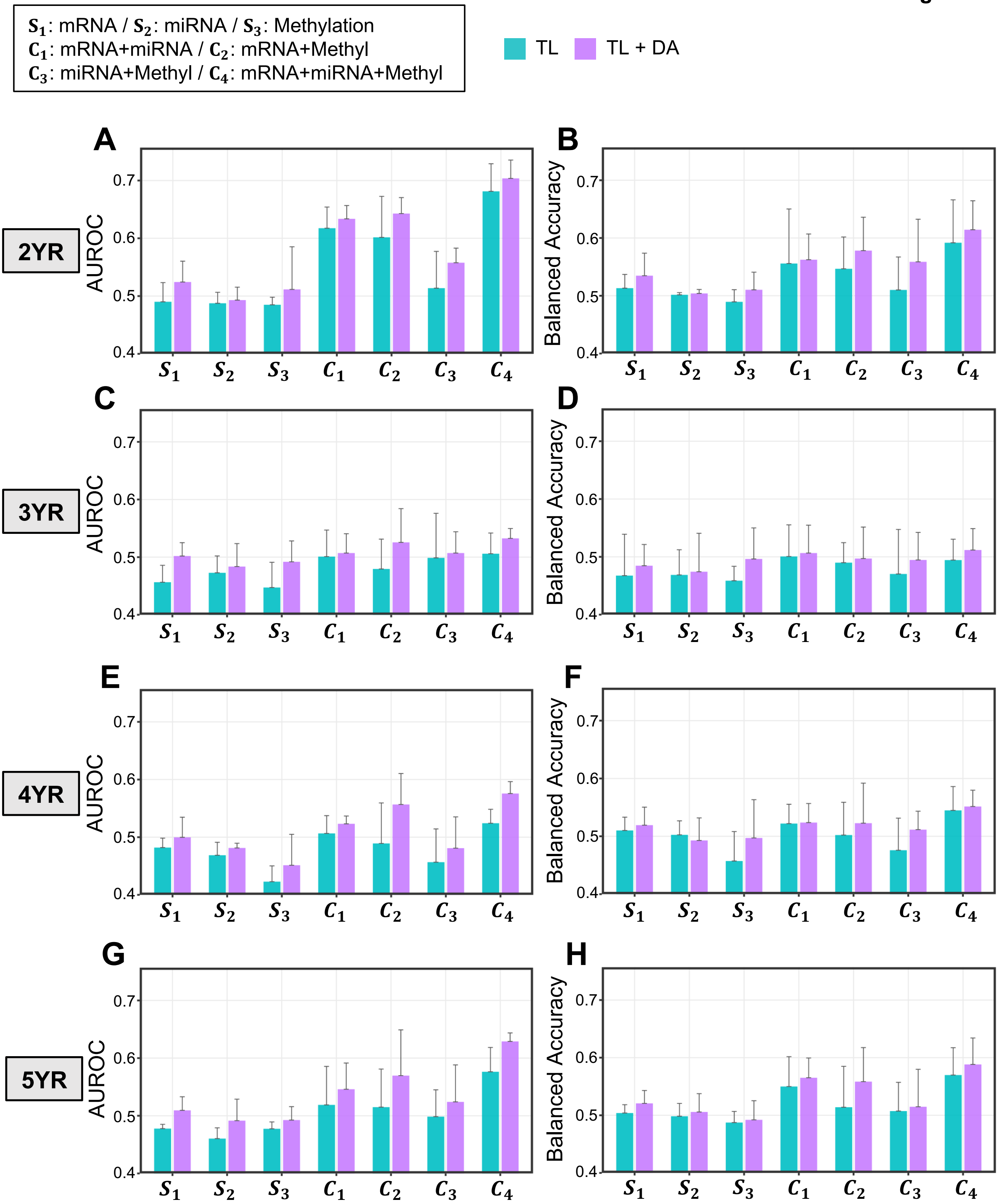

$S_1$: mRNA / $S_2$: miRNA / $S_3$: Methylation
$C_1$: mRNA+miRNA / $C_2$: mRNA+Methyl
$C_3$: miRNA+Methyl / $C_4$: mRNA+miRNA+Methyl
TL
TL + DA
A
B
C
D
E
F
G
H
2YR
3YR
4YR
5YR
AUROC
Balanced Accuracy
0.4
0.5
0.6
0.7
$S_1$ $S_2$ $S_3$ $C_1$ $C_2$ $C_3$ $C_4$

**Figure S2. Comparison of prediction performance between transfer learning and MOTLAB across single-, two-, and three-omics combinations.**

(**A-H**) Prediction performance of conventional transfer learning and MOTLAB was evaluated using single-, two-, and three-omics data across four progression-free interval (PFI) predictions (2-, 3-, 4-, and 5-year). The left column shows the area under the receiver operating characteristic curve (AUROC), and the right column shows balanced accuracy (BACC). Rows correspond to the 2-year (**A-B**), 3-year (**C-D**), 4-year (**E-F**), and 5-year (**G-H**) PFI prediction models. Performance was evaluated separately for each omics combination, including mRNA ($\boldsymbol{S_1}$), miRNA ($\boldsymbol{S_2}$), Methylation ($\boldsymbol{S_3}$), mRNA + miRNA ($\boldsymbol{C_1}$), mRNA + Methylation ($\boldsymbol{C_2}$), miRNA + Methylation ($\boldsymbol{C_3}$), and mRNA + miRNA + Methylation ($\boldsymbol{C_4}$). Bars represent the mean performance across repeated random seeds, and error bars indicate one-sided upper standard deviations.

**Figure S3**

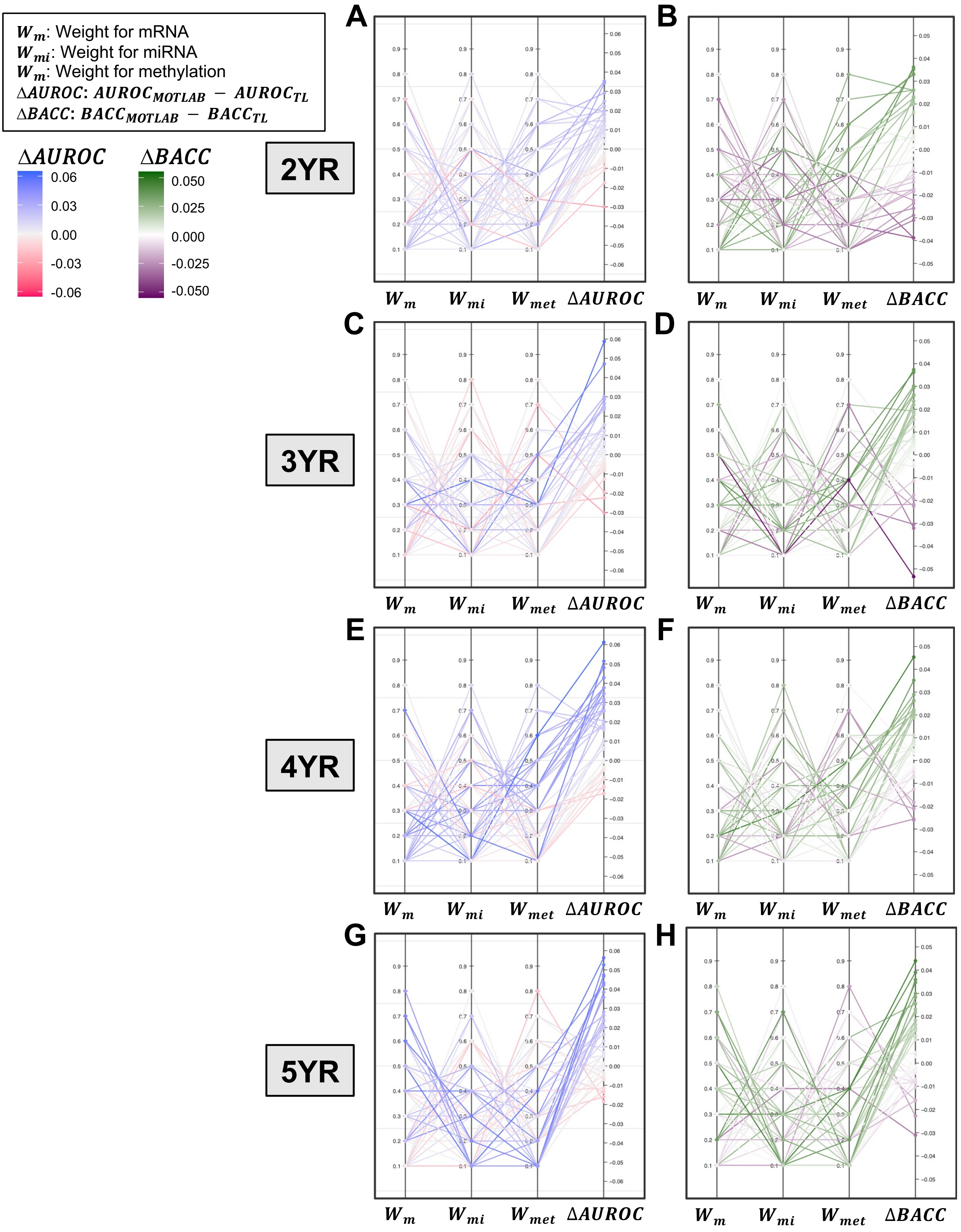
$W_m$: Weight for mRNA
$W_{mi}$: Weight for miRNA
$W_m$: Weight for methylation
$\Delta AUROC$: $AUROC_{MOTLAB} - AUROC_{TL}$
$\Delta BACC$: $BACC_{MOTLAB} - BACC_{TL}$
$\Delta AUROC$
0.06
0.03
0.00
-0.03
-0.06
$\Delta BACC$
0.050
0.025
0.000
-0.025
-0.050
2YR
3YR
4YR
5YR
A
B
C
D
E
F
G
H
$W_m$
$W_{mi}$
$W_{met}$
$\Delta AUROC$
$\Delta BACC$

**Figure S3. Three-omics weighting schemes and performance differences between MOTLAB and transfer learning alone.**

(**A–H**) Parallel-coordinate plots showing predefined three-omics weighting schemes for mRNA, miRNA, and DNA methylation and their corresponding performance differences between MOTLAB and transfer learning alone. ΔAUROC was defined as AUROC_MOTLAB − AUROC_TL, and ΔBACC was defined as BACC_MOTLAB − BACC_TL. Results are shown for 2-year PFI prediction using ΔAUROC (**A**) and ΔBACC (**B**), 3-year PFI prediction using ΔAUROC (**C**) and ΔBACC (**D**), 4-year PFI prediction using ΔAUROC (**E**) and ΔBACC (**F**), and 5-year PFI prediction using ΔAUROC (**G**) and ΔBACC (**H**). First three lines represent weights for individual omics data. Colors indicate the magnitude and direction of performance difference relative to transfer learning alone with last line. Positive ΔAUROC or ΔBACC values indicate higher performance for MOTLAB, whereas negative values indicate lower performance for MOTLAB. Grid searching for optimizing weights was used to assess whether MOTLAB performance depended on how weights were distributed across the three omics modalities.

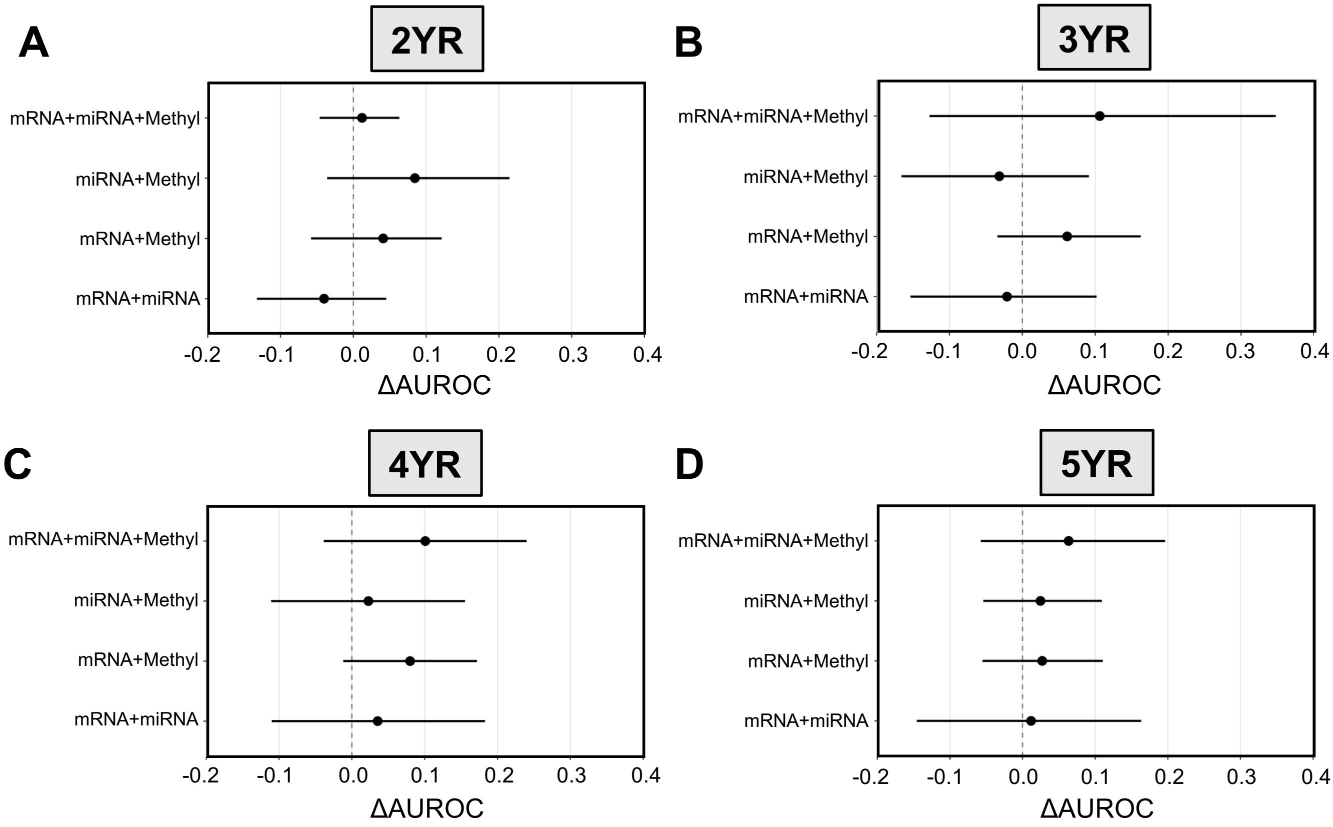


**Figure S4. Patient-level paired bootstrap estimates of the AUROC difference between MOTLAB and conventional transfer learning across PFI prediction settings.**
(**A–D**) Patient-level paired bootstrap estimates of the difference in AUROC for predicting progression-free interval (PFI) at 2-year (**A**), 3-year (**B**), 4-year (**C**), and 5-year (**D**) PFI predictions. For each omics combination, the point represents the observed ΔAUROC calculated from patient-level mean prediction scores after averaging across matched random seeds shared between MOTLAB and conventional transfer learning. Horizontal lines indicate bias-corrected and accelerated 95% confidence intervals estimated from 10,000 outcome-stratified paired bootstrap resamples. The vertical dashed line denotes ΔAUROC = 0 (no performance difference between models). Positive ΔAUROC values indicate higher discriminative performance of MOTLAB relative to conventional transfer learning.

Figure S5

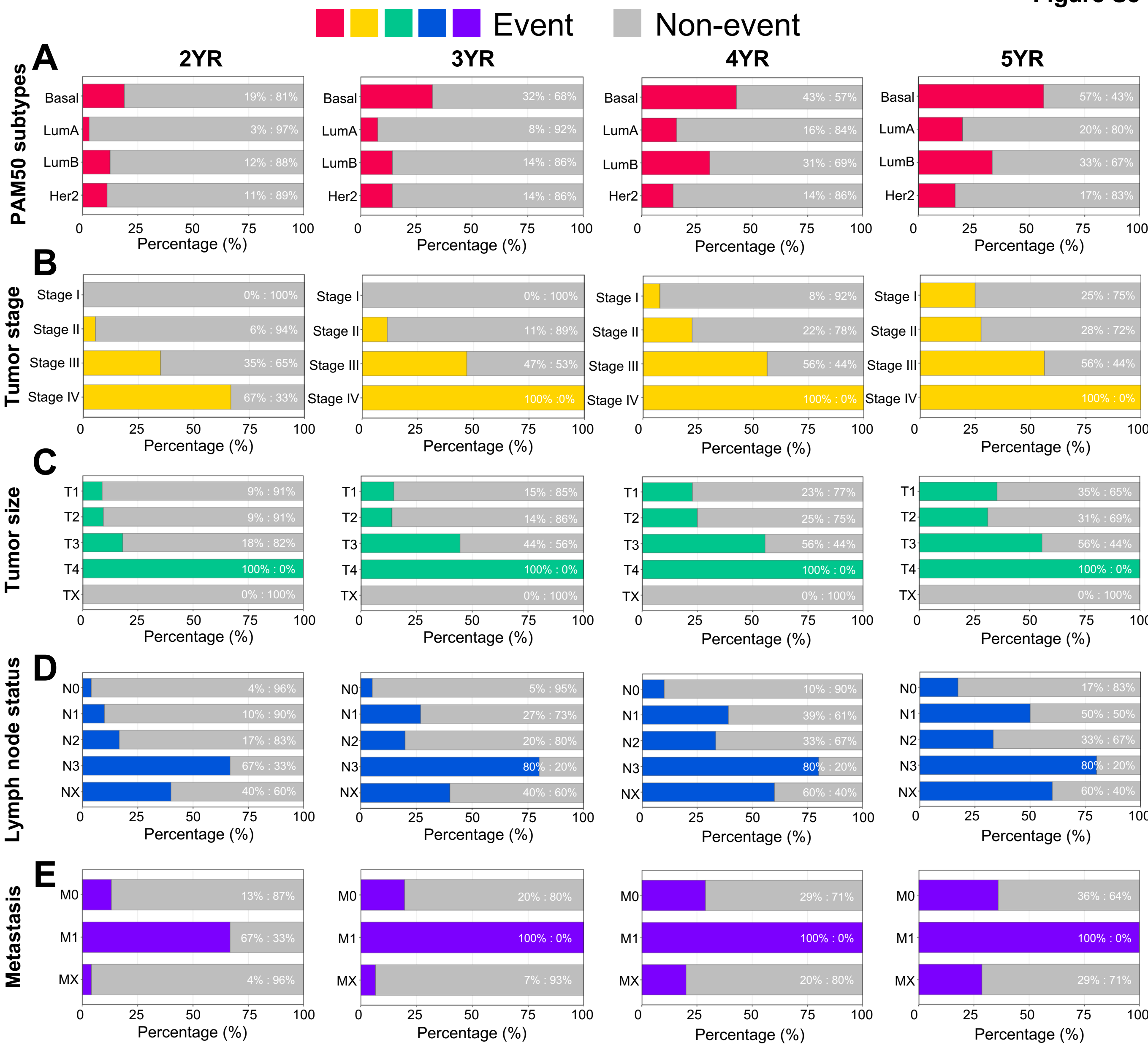


**Figure S5. Event and non-event composition across molecular and clinicopathologic subgroups.**

(**A–E**) Subgroup-level distributions of PFI events and non-events across 2-, 3-, 4-, and 5-year PFI prediction settings. Event and non-event proportions are shown for PAM50 intrinsic subtype groups (**A**), tumor stage groups (**B**), tumor size groups (**C**), lymph node status groups (**D**), and metastasis status groups (**E**). Each horizontal bar is scaled to 100% within a given subgroup level and PFI prediction setting. Colored segments indicate PFI events by the corresponding PFI prediction setting, and gray segments indicate non-events or favorable status by that PFI prediction setting. Percent labels indicate within-subgroup event and non-event proportions. These plots provide subgroup-level outcome-composition context for interpreting subgroup performance estimates in Figure 5.

### Figure S6

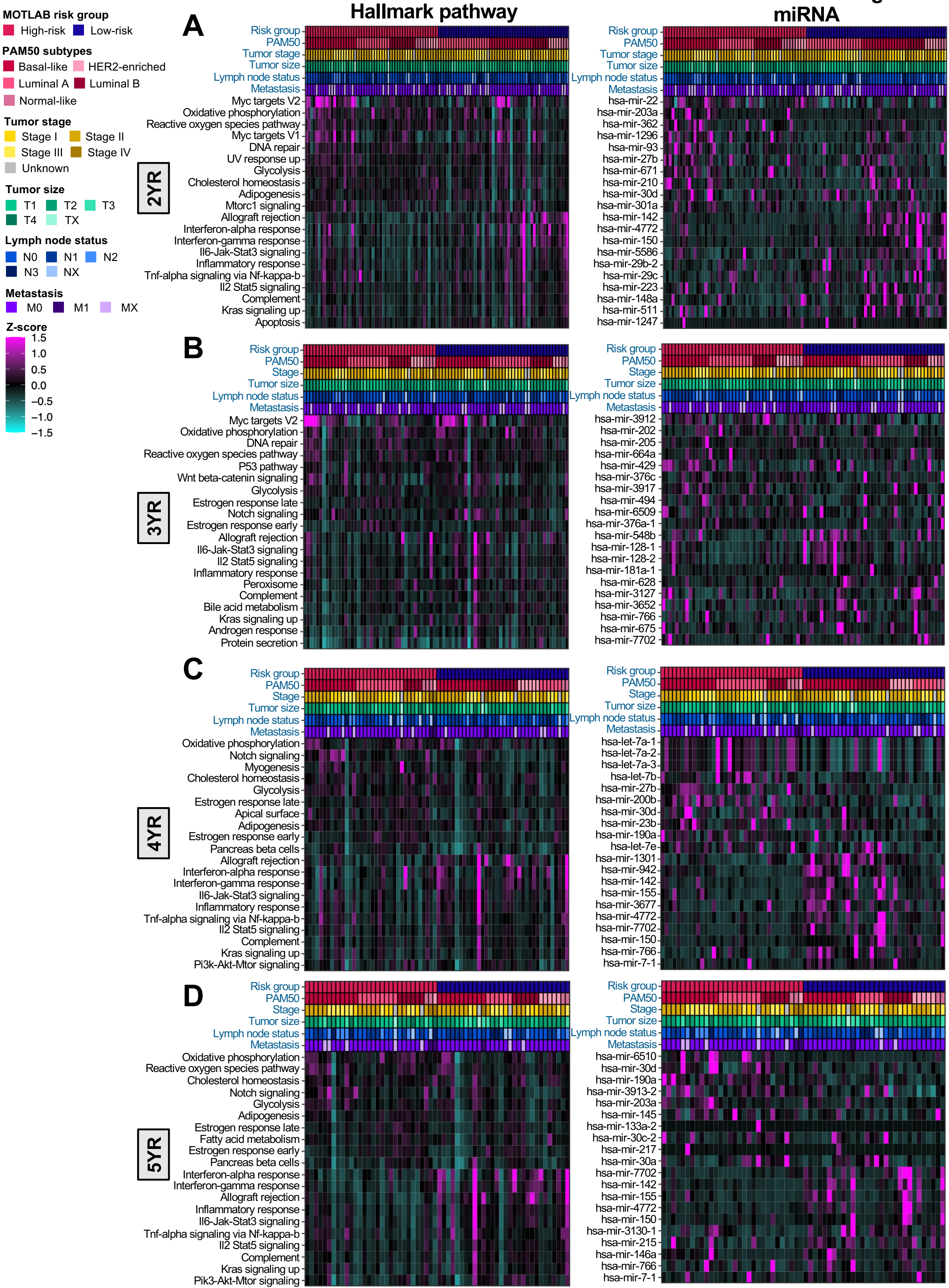

Hallmark pathway
miRNA
MOTLAB risk group
High-risk
Low-risk
PAM50 subtypes
Basal-like
HER2-enriched
Luminal A
Luminal B
Normal-like
Tumor stage
Stage I
Stage II
Stage III
Stage IV
Unknown
Tumor size
T1
T2
T3
T4
TX
Lymph node status
N0
N1
N2
N3
NX
Metastasis
M0
M1
MX
Z-score
1.5
1.0
0.5
0.0
−0.5
−1.0
−1.5
A
2YR
Risk group
PAM50
Tumor stage
Tumor size
Lymph node status
Metastasis
Myc targets V2
Oxidative phosphorylation
Reactive oxygen species pathway
Myc targets V1
DNA repair
UV response up
Glycolysis
Cholesterol homeostasis
Adipogenesis
Mtorc1 signaling
Allograft rejection
Interferon-alpha response
Interferon-gamma response
Il6-Jak-Stat3 signaling
Inflammatory response
Tnf-alpha signaling via Nf-kappa-b
Il2 Stat5 signaling
Complement
Kras signaling up
Apoptosis
hsa-mir-22
hsa-mir-203a
hsa-mir-362
hsa-mir-1296
hsa-mir-93
hsa-mir-27b
hsa-mir-671
hsa-mir-210
hsa-mir-30d
hsa-mir-301a
hsa-mir-142
hsa-mir-4772
hsa-mir-150
hsa-mir-5586
hsa-mir-29b-2
hsa-mir-29c
hsa-mir-223
hsa-mir-148a
hsa-mir-511
hsa-mir-1247
B
3YR
Risk group
PAM50
Stage
Tumor size
Lymph node status
Metastasis
Myc targets V2
Oxidative phosphorylation
DNA repair
Reactive oxygen species pathway
P53 pathway
Wnt beta-catenin signaling
Glycolysis
Estrogen response late
Notch signaling
Estrogen response early
Allograft rejection
Il6-Jak-Stat3 signaling
Il2 Stat5 signaling
Inflammatory response
Peroxisome
Complement
Bile acid metabolism
Kras signaling up
Androgen response
Protein secretion
hsa-mir-3912
hsa-mir-202
hsa-mir-205
hsa-mir-664a
hsa-mir-429
hsa-mir-376c
hsa-mir-3917
hsa-mir-494
hsa-mir-6509
hsa-mir-376a-1
hsa-mir-548b
hsa-mir-128-1
hsa-mir-128-2
hsa-mir-181a-1
hsa-mir-628
hsa-mir-3127
hsa-mir-3652
hsa-mir-766
hsa-mir-675
hsa-mir-7702
C
4YR
Risk group
PAM50
Stage
Tumor size
Lymph node status
Metastasis
Oxidative phosphorylation
Notch signaling
Myogenesis
Cholesterol homeostasis
Glycolysis
Estrogen response late
Apical surface
Adipogenesis
Estrogen response early
Pancreas beta cells
Allograft rejection
Interferon-alpha response
Interferon-gamma response
Il6-Jak-Stat3 signaling
Inflammatory response
Tnf-alpha signaling via Nf-kappa-b
Il2 Stat5 signaling
Complement
Kras signaling up
Pi3k-Akt-Mtor signaling
hsa-let-7a-1
hsa-let-7a-2
hsa-let-7a-3
hsa-let-7b
hsa-mir-27b
hsa-mir-200b
hsa-mir-30d
hsa-mir-23b
hsa-mir-190a
hsa-let-7e
hsa-mir-1301
hsa-mir-942
hsa-mir-142
hsa-mir-155
hsa-mir-3677
hsa-mir-4772
hsa-mir-7702
hsa-mir-150
hsa-mir-766
hsa-mir-7-1
D
5YR
Risk group
PAM50
Stage
Tumor size
Lymph node status
Metastasis
Oxidative phosphorylation
Reactive oxygen species pathway
Cholesterol homeostasis
Notch signaling
Glycolysis
Adipogenesis
Estrogen response late
Fatty acid metabolism
Estrogen response early
Pancreas beta cells
Interferon-alpha response
Interferon-gamma response
Allograft rejection
Inflammatory response
Il6-Jak-Stat3 signaling
Tnf-alpha signaling via Nf-kappa-b
Il2 Stat5 signaling
Complement
Kras signaling up
Pik3-Akt-Mtor signaling
hsa-mir-6510
hsa-mir-30d
hsa-mir-190a
hsa-mir-3913-2
hsa-mir-203a
hsa-mir-145
hsa-mir-133a-2
hsa-mir-30c-2
hsa-mir-217
hsa-mir-30a
hsa-mir-7702
hsa-mir-142
hsa-mir-155
hsa-mir-4772
hsa-mir-150
hsa-mir-3130-1
hsa-mir-215
hsa-mir-146a
hsa-mir-766
hsa-mir-7-1

**Figure S6. Patient-level Hallmark pathway and miRNA expression heatmaps associated with MOTLAB risk groups.**

(**A–D**) Patient-level heatmaps of Hallmark pathway scores and miRNA abundance across 2-year (**A**), 3-year (**B**), 4-year (**C**), and 5-year (**D**) PFI prediction settings. The left heatmap in each PFI prediction setting shows selected MSigDB Hallmark scores, and the right heatmap shows selected miRNA abundance features corresponding to the molecular associations summarized in **Fig 7**. Rows show the top 10 positively and top 10 negatively correlated Hallmark scores or miRNA features with MOTLAB risk scores based on Spearman's ρ. Columns represent individual patients ordered by MOTLAB high-risk group first, followed by MOTLAB low-risk group; within each risk group, patients were further ordered by subgroup-level sample size and MOTLAB risk scores. Annotation tracks show MOTLAB risk group, PAM50 intrinsic subtype, tumor stage, tumor size, lymph node status, and metastasis status. Although TCGA also classified a subset of samples as the Normal-like PAM50 subtype, these samples were excluded in the PAM50 subtype analysis but are retained in other subtypes analysis. PFI event status was not included as a heatmap annotation. Hallmark scores were calculated as the mean of gene-wise z-scored mRNA expression values for genes present in each MSigDB Hallmark set, rather than by GSVA or ssGSEA. miRNA values were z-scored within each miRNA feature. Heatmap colors represent row-wise scaled values, capped at ±1.5 for Hallmark scores and ±3 for miRNA abundance.

**Figure S7**

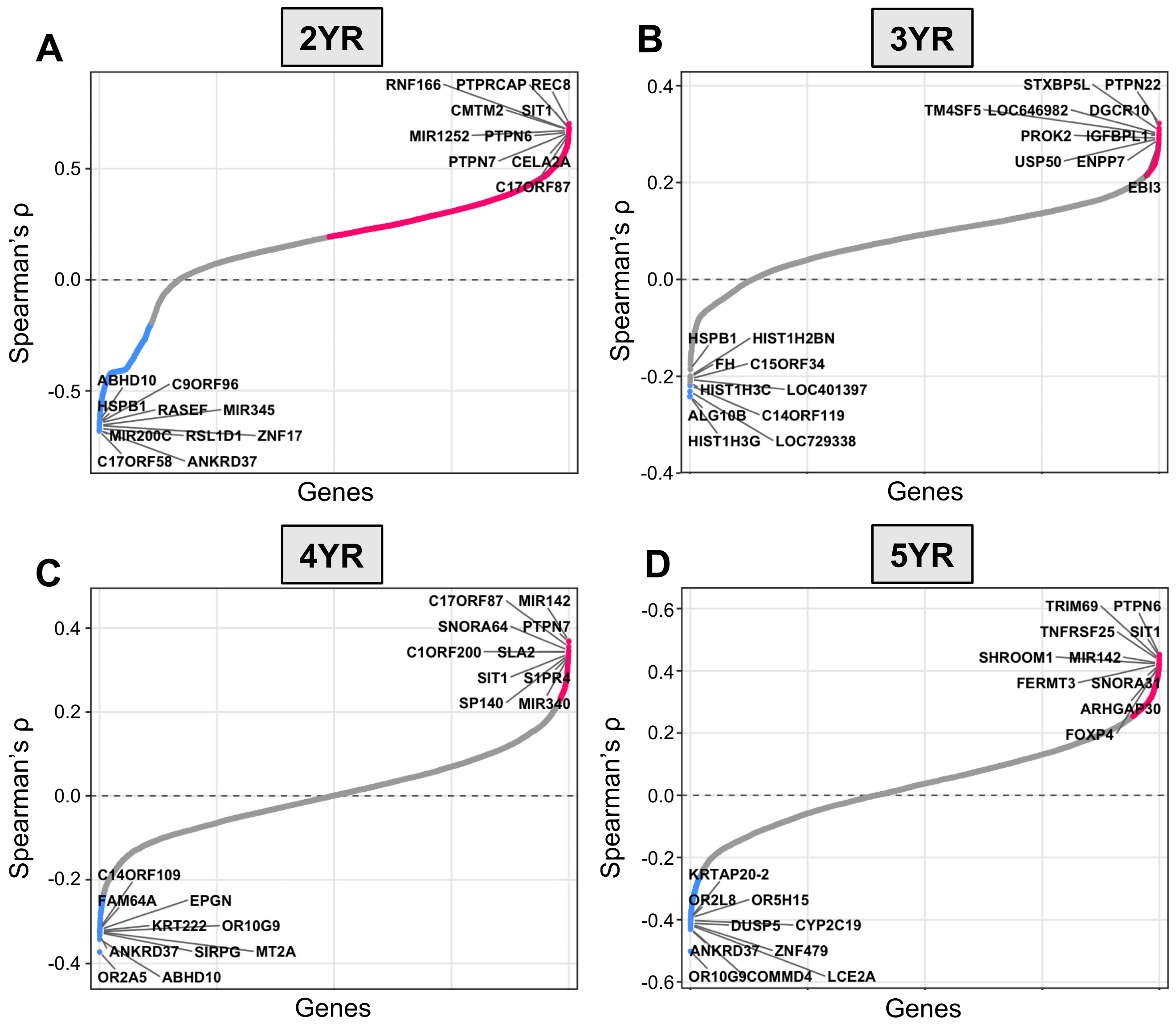


**Figure S7. Gene-level DNA methylation correlations with MOTLAB risk scores across PFI prediction settings.**

(**A–D**) Ranked gene-level DNA methylation Spearman correlations with MOTLAB risk scores across 2-year (**A**), 3-year (**B**), 4-year (**C**), and 5-year (**D**) PFI prediction settings. Probe-level methylation features were summarized at the gene level by averaging available methylation values mapped to each gene. Each point represents one gene-level methylation summary ranked by Spearman's ρ with MOTLAB risk. Colored points indicate positive and negative top10 gene-level methylations, and gray points indicate nominal raw $p \geq 0.05$. Positive ρ values indicate higher methylation with increasing MOTLAB risk, whereas negative ρ values indicate lower methylation with increasing MOTLAB risk. Labeled genes represent the 10 highest-ρ and 10 lowest-ρ genes selected for visualization. The horizontal dashed line indicates Spearman's ρ = 0.
